# microRNA mir-598-3p mediates susceptibility to stress-enhancement of remote fear memory

**DOI:** 10.1101/584326

**Authors:** Meghan E Jones, Stephanie E. Sillivan, Sarah Jamieson, Gavin Rumbaugh, Courtney A. Miller

## Abstract

microRNAs (miRNAs) have emerged as potent regulators of learning, recent memory and extinction. However, our understanding of miRNAs directly involved in regulating complex psychiatric conditions perpetuated by aberrant memory, such as in posttraumatic stress disorder (PTSD), remains limited. To begin to address the role of miRNAs in persistent memory, we performed small-RNA sequencing on basolateral amygdala (BLA) tissue to identify miRNAs altered by auditory fear conditioning (FC) one month after training. mir-598-3p, a highly conserved miRNA previously unstudied in the brain, was downregulated in the BLA. Further decreasing BLA mir-598-3p levels did not alter the expression or extinction of the remote fear memory. Given that stress is a critical component in PTSD, we next assessed the impact of stress-enhanced fear learning (SEFL) on mir-598-3p levels, finding the miRNA is elevated in the BLA of male, but not female, mice susceptible to the effects of stress in SEFL. Accordingly, intra-BLA inhibition of mir-598-3p interfered with expression and extinction of the remote fear memory in male, but not female, mice. This effect could not be attributed to an anxiolytic effect of miRNA inhibition. Finally, bioinformatic analysis following quantitative proteomics on BLA tissue collected 30 days post-SEFL training identified putative mir-598-3p targets and related pathways mediating the differential susceptibility, with evidence for regulation of the actin cytoskeleton.

## INTRODUCTION

Persistent, maladaptive long-term memories are an integral part of several psychiatric disorders, including substance use disorder (SUD, [1]) and post-traumatic stress disorder (PTSD, [2]). While much of the current literature has focused on mechanisms supporting recent memory acquisition and consolidation [3], a greater understanding of the neurobiological mechanisms governing persistent, enhanced remote memory could provide new insight into treatment of memory-related disorders. Additionally, memories that drive unwanted flashbacks of trauma in PTSD are unique from normal episodic memories in that they are intrusive and more life-like in nature [4]. Stress, even one acute episode, can impose lasting changes on the brain [5-7]. But how stress-induced changes influence or interact with remote memory mechanisms is not well-characterized. Here, we examined the role of microRNAs (miRNAs) in the persistence of a remote fear memory with and without exposure to acute stress.

miRNAs are small, ∼ 22-24 nucleotide non-coding RNAs that repress translation of target mRNAs [8, 9]. miRNAs interact with multiple functional targets directly at the 3’ untranslated region of the target mRNA based on complimentary sequences and, therefore, are capable of supporting the complex patterns of gene regulation required to support persistent, remote memory. In an early example linking miRNAs to recent memory, knockdown of Dicer1, an enzyme required for mature miRNA processing, enhanced memory performance in the Morris Water Maze, trace fear conditioning, and contextual fear conditioning [10], confirming that miRNA function influences memory performance in several different types of tasks. More specifically, Gao and colleagues demonstrated that hippocampal miR-134 overexpression impairs memory acquisition by targeting both cAMP response binding element protein (CREB) and brain-derived neurotrophic factor (BDNF) in a contextual fear conditioning task [11]. We have previously shown that cued fear conditioning downregulates miR-182 in the lateral amygdala and that overexpression of miR-182 in this region during training disrupted expression of the fear memory the following day [12]. While these reports and others provide evidence that miRNAs can influence memory formation [13-15], studies to date have not addressed their impact on remote memory. One goal of the experiments reported here is to gain a better understanding of the role of miRNAs in the persistence of remote memory. We focused our analyses on the basolateral amygdala (BLA) because of the known contribution of this region to fear learning and memory storage at remote time points [16-19]

We recently demonstrated that exposure to a single acute stressor (120 minutes of restraint stress) induces a change in the molecular profile of the BLA, including differential expression of 18 miRNAs an entire month after stress [5]. Differential miRNA expression has also been reported in the serum of patients with PTSD [20] [21] and between rats selectively bred for high or low stress reactivity [22]. The behavioral consequences of stress exposure have also been linked to miRNA function in the amygdala in several reports. For example, Haramati and colleagues identified several miRNAs regulated by acute stress and showed that central amygdala (CeA) overexpression of mir-34c protected against stress-induced increases in anxiety-like behavior in the light/dark transfer test [23]. Similarly, transgenic overexpression of mir-26a-2 protected against increases in anxiety-like behavior induced by social defeat stress [24]. Thus, miRNAs regulate both stress-induced behaviors and learning and memory, suggesting they may make an important contribution to PTSD susceptibility.

## RESULTS

### Role of amygdala mir-598-3p in remote fear memory

We employed small RNA sequencing (smRNA-seq) to identify miRNAs in the BLA that may contribute to the support of a remote fear memory. To isolate memory-associated miRNAs, mice that underwent auditory FC were compared to naïve mice and those exposed to an unpaired (UNP) protocol. The UNP protocol consisted of the same number of tone and shock presentations, but the tone did not coterminate with the foot shock. Tissue from the BLA was collected 30 days after training and small-RNA libraries were prepared and sequenced from biological replicate samples. This was performed at two independent sequencing facilities on two independent animal cohorts and the data from each was compared to identify miRNAs differentially regulated in both runs (**Figure 1A**). To confirm that FC, but not the UNP protocol, resulted in cue-specific learning, a separate cohort of mice underwent a retrieval session in a distinct context 30 days after training. As expected, FC but not UNP mice displayed remote fear memory expression (**Figure 1B;** RM ANOVA, main effect of group: F(1,65) = 9.991, *p* = 0.002).

**Figure 1.**
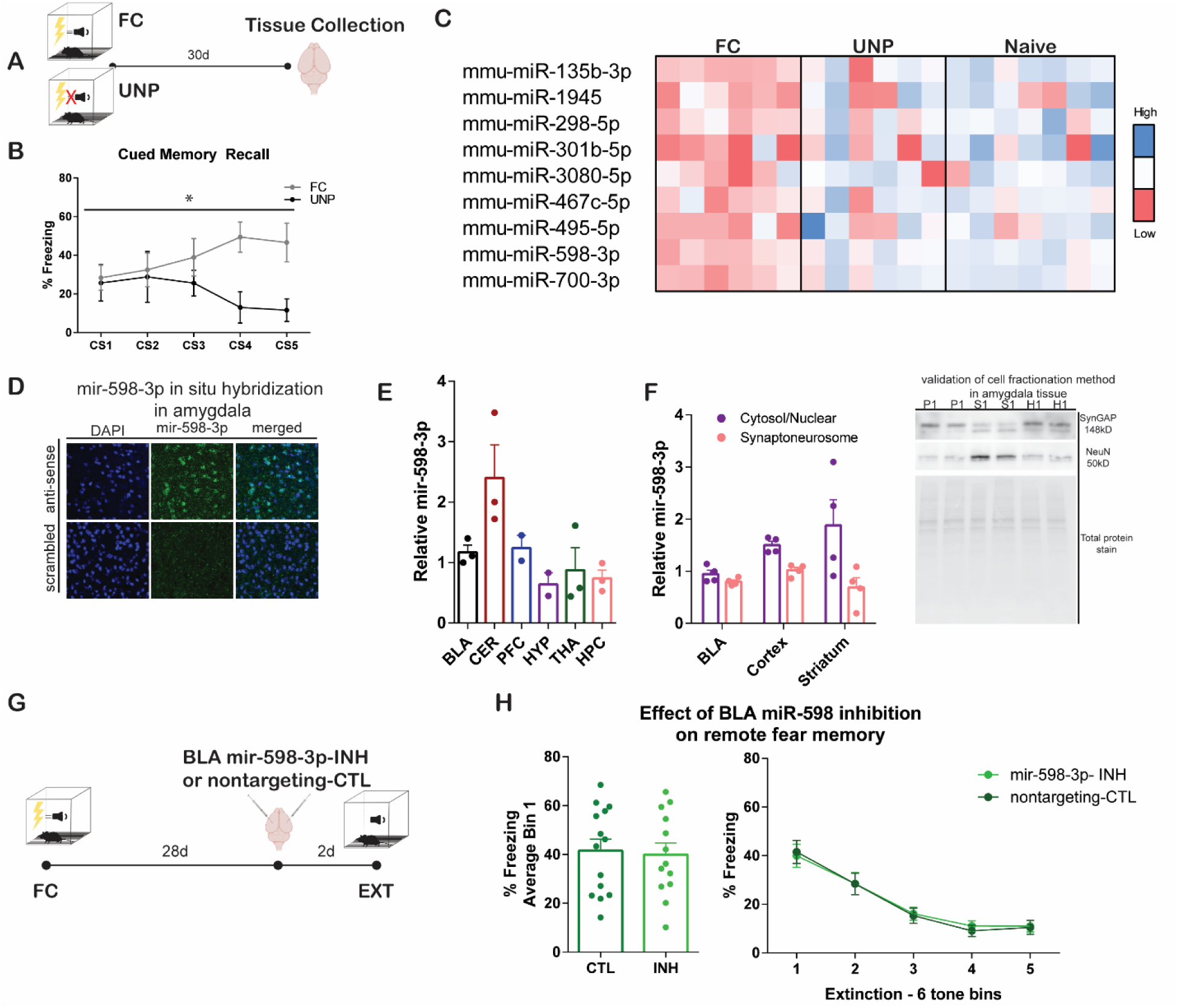
Role of mir-598-3p in remote fear memory expression. (A) Schematic of experimental design for smRNA-seq experiments. (B) Training resulted cue-specific memory expression in FC mice, but not UNP mice, * main effect of group, *p* < 0.05. (C) miRNAs identified as differentially expressed by 0.5 log_2_ fold change between both FC and UNP and FC and naïve comparisons. (D) BLA expression of mir-598-3p by fluorescent in situ hybridization. (E) mir-598-3p relative expression in BLA, cerebellum (CER), prefrontal cortex (PFC), hypothalamus (HYP), thalamus (THA), and hippocampus (HPC). Values are relative to the average ddCt value of all regions. (F) Cell fractionation protocol used to measure relative mir-598-3p in the synaptoneurosome and cytosol in the BLA, cortex and striatum (STR). Western blotting confirmed the pellet (P) expressed a known synaptic marker (SynGAP) while the supernatant (S) expressed a known nuclear marker (NeuN), validating the cell fractionation method to isolate the synaptoneurosome and cytosol. (G) Schematic shows mir-598-3p inhibition in the BLA 28 days after cued FC, followed by a test of remote fear memory expression during an extinction session in a unique context. (H) mir-598-3p expression did not alter fear memory recall or extinction. FC n = 13-14 per group. Data presented are % freezing normalized to individual pretone freezing in 6 tone bins and as average freezing during the first 6 tone bin (memory recall).

miRNAs that were identified as differentially expressed in both smRNA-seq runs by at least 0.5 log2 fold change between FC mice and both UNP and naïve are displayed in **Figure 1C** (and **Supplemental File 1**). Nine miRNAs were differentially expressed, all of which were downregulated in the FC condition. mir-598-3p was selected for further analysis based on its novelty, as its function had not yet been explored in the brain. After confirming a decrease in mir-598-3p levels one month after FC via qPCR (Naïve: mean = 1.0 ± 0.15; FC: mean = 0.66 ± 0.08; one-tailed t-test: t_(17)_ = 2.181, *p* = 0.022), we examined its expression pattern in the BLA by fluorescent in situ hybridization, finding it broadly expressed throughout the region (**Figure 1D**). Further, we characterized its relative values in the BLA, cerebellum (CER), prefrontal cortex (PFC), hypothalamus (HYP), thalamus (THA), and hippocampus (HPC) of naïve mice (**Figure 1E**), finding the highest expression in CER and lowest in HYP. Cellular localization analysis revealed that mir-598-3p is expressed throughout cells, including in the synaptoneurosome fraction, in the BLA, cortex, and striatum (**Figure 1F**). Interestingly, the ratio of mir-598-3p expression in the synaptoneurosome relative to the rest of the cell was highest in the BLA.

We next determined the potential for in vivo inhibition of mir-598-3p within the BLA to enhance remote fear memory expression. Mice underwent cued fear conditioning and received either a synthetic hairpin miRNA inhibitor targeting mir-598-3p (mir-598-3p-INH) or a control inhibitor 28 days later (**Figure 1G**). Remote fear memory was assessed based on freezing during the first set of tone presentations (Bin 1) in a distinct context, followed by assessment of extinction in the same session with the presentation of an additional 24 tones. There was no effect on retrieval, as freezing was equivalent during Bin 1 tone presentations (**Figure 1H;** two-tailed independent samples t-test: t_(25)_ = 0.240, *p* = 0.812). BLA mir-598-3p inhibition also did not alter remote fear memory extinction (**Figure 1H**). We observed a significant effect of time (two-way RM ANOVA, main effect of tone: F_(4,125)_ = 27.642, *p* < 0.001), indicating that freezing behavior diminished across the extinction session, but no effect of mir-598-3p-INH treatment (F_(1,125)_ = 0.024, *p* = 0.878). There was also no effect of treatment on the rate of extinction across the session (two-tailed independent samples t-test: t_(25)_ =0.701, *p* =0.490).

### Amygdala mir-598-3p is differentially expressed in stress susceptible and resilient mice and regulates stress-enhanced fear memory (SEFM)

We previously developed a protocol through which stress prior to FC results in an enhancement of fear memory in a subgroup of mice [25]. Specifically, male mice display differential susceptibility to the effects of stress and the susceptible (SS) and resilient (SR) animals can be identified by their freezing behavior during the final minute of FC, without additional phenotyping and prior to remote memory recall. Expression of the remote SEFM is accompanied by immediate early gene activation in the BLA and changes to BLA mRNA expression of genes reported to be altered in subjects diagnosed with PTSD, including adenylate cyclase activating polypeptide 1 (PACAP), BDNF, and tyrosine hydroxylase. In contrast to males, female mice display a consistent susceptible-like phenotype following SEFL, exhibiting SEFM comparable to that of SS males [25].

Given the behavioral interaction between stress and FC and the involvement of the BLA, we next assessed the long-term impact of restraint stress alone or SEFL training on BLA mir-598-3p levels. BLA tissue was collected for qPCR analysis of mir-598-3p 30 days after stress, FC, or SEFL and freezing behavior was analyzed during the last minute of training in SEFL males to identify SS and SR subgroups (**Figure 2A**; [25]). In males, mir-598-3p was differentially expressed across the five groups tested one month after behavioral manipulation (**Figure 2B**, left panel; one-way ANOVA: F_(4,36)_ = 9.447, *p* < 0.001). LSD post hoc comparisons indicated that stress increased mir-598-3p (*p* = 0.004), while FC decreased the miRNA, (*p* = 0.043), relative to naïve mice. Among mice that underwent SEFL, SS mice exhibited significantly higher mir-598-3p than SR mice (*p* = 0.044). Interestingly, female mice exhibited no change in mir-598-3p after stress, FC, or SEFL (**Figure 2B**, right panel; one-way ANOVA: F_(3,22)_ = 0.123, *p* = 0.945).

**Figure 2.**
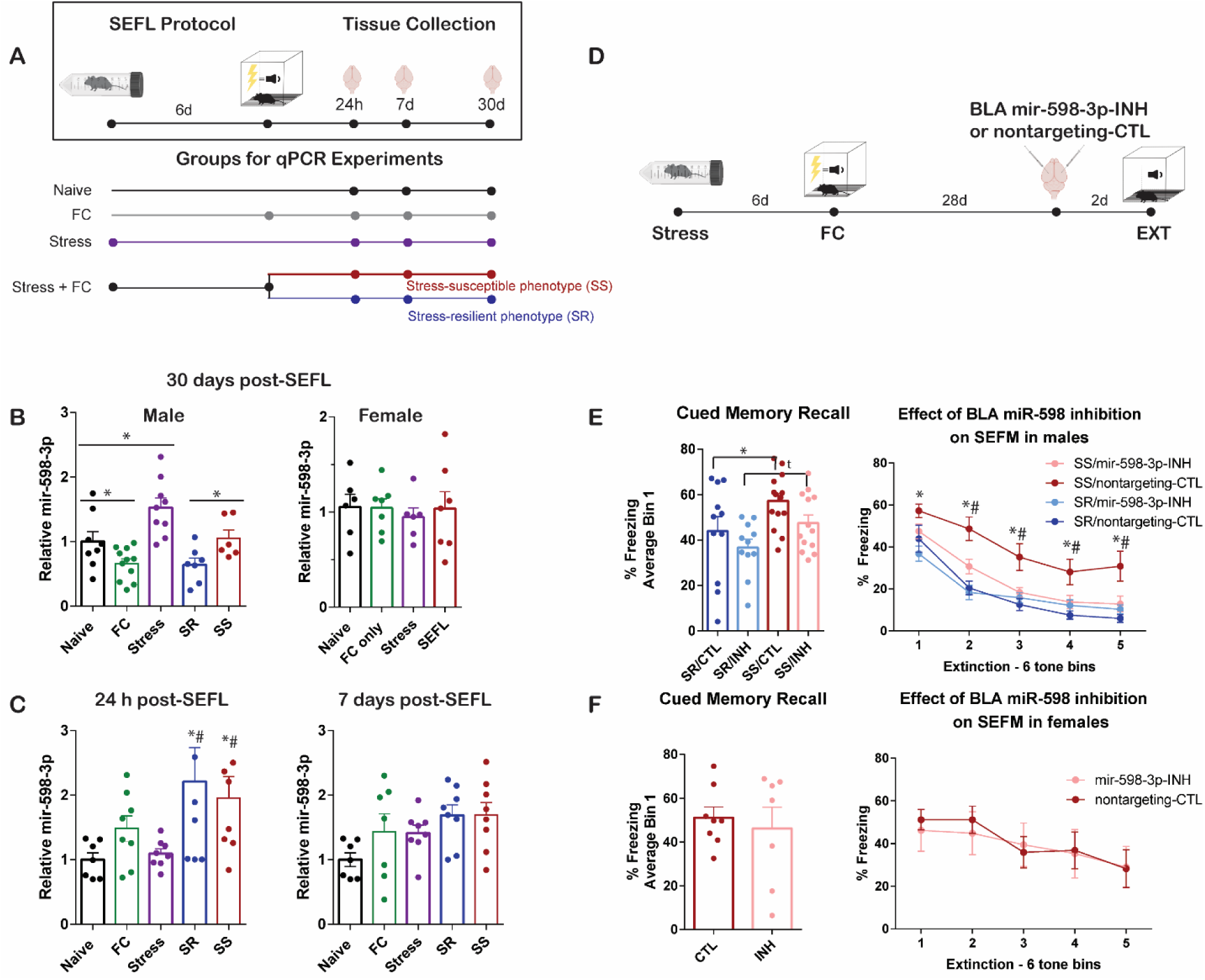
Amygdala mir-598-3p is differentially expressed in stress susceptible and resilient mice and regulates stress-enhanced fear memory (SEFM) in susceptible mice. (A) Schematic of experimental design. (B) Relative mir-598-3p expression in males and females 30 days after FC, Stress, or SEFL. Male n = 7-11 per group, Female n = 6-7 per group; * = *p* < 0.05. (C) Relative mir-598-3p expression in males 24 hours and 7 days after FC, Stress, or SEFL. 24h n = 6-8, 7d n = 7-8 per group; * *p* < 0.05 vs. naïve, # *p* < 0.05 vs. stress. (D). Schematic of mir-598-3p inhibition experimental design. (E) BLA mir-598-3p inhibition attenuated stress-enhanced fear memory (SEFM) in male SS mice. SR n = 11 per group, SS n = 13-14 per group; * SR/CTL vs. SS/CTL *p* < 0.05; # SS/CTL vs. SS/INH *p* < 0.05; *t p* = 0.052. (F) BLA mir-598-3p inhibition did not alter SEFM in female mice. Female n = 7-8 per group. Behavioral data are presented as % freezing normalized to individual pretone freezing in 6 tone bins and as average freezing during the first 6 tone bin (memory recall).

To determine the time course leading up to the mir-598-3p expression differences observed in males 30 days after training, we ran additional cohorts, collecting tissue 24 hours or 7 days later (**Figure 2A**). The results indicate a surprisingly dynamic level of regulation of mir-598-3p in the BLA over time. At 24 hours, mir-598-3p was differentially expressed (**Figure 2C**, left panel; one-way ANOVA: F_(4,34)_ = 3.016, *p* = 0.031). LSD post hoc comparisons revealed that mir-598-3p was significantly increased in SEFL mice, relative to naïve and stress alone, and that the difference was significant for both SS (naïve *p* = 0.009, stress *p* = 0.049) and SR subgroups (naïve *p* = 0.036, stress *p* = 0.012). At 7 days, there were no significant group differences in mir-598-3p expression (**Figure 2C**, right panel; one-way ANOVA: F_(4,33)_ = 2.371, *p* = 0.072), yet this continues to change, as evidenced by the 30 day data (**Figure 2B**). These results demonstrate that SEFL induces an initial increase in BLA mir-598-3p that persists over time in SS mice, while SR levels drop to FC only control levels by 30 days. One interpretation is that the stress-induced increase in mir-598-3p dominates the response to SEFL in SS mice, while the FC-induced decrease in mir-598-3p dominates the response to SEFL in SR mice over time. Subsequently, we hypothesized that inhibition of mir-598-3p in SS mice would protect against the development of SEFM, mimicking an SR-like memory phenotype.

We next determined the effect of inhibiting mir-598-3p on remote SEFM in the BLA one month after SEFL training. Twenty-eight days after SEFL training, male and female mice received bilateral intra-BLA infusions of mir-598-3p-INH or a nontargeting control and remote fear memory was assessed two days later (**Figure 2D**). During the first Bin of tone presentations, SS males froze more than SR males (F_(1,45)_ = 8.061, *p* = 0.007) and there was a strong trend towards a main effect of treatment (F_(1,45)_ = 4.0, *p* = 0.052), indicating an attenuation of remote fear memory expression with mir-598-3p inhibition. Further, freezing behavior diminished across the entire session (**Figure 2E**; three-way RM ANOVA, main effect of tone: F_(4,225)_ = 36.515, *p* < 0.001) and SS control mice exhibited the expected SEFM, in that they displayed higher freezing than SR controls (main effect of population: F_(1,225)_ = 51.349, *p* <0.001). Consistent with our hypothesis, BLA inhibition of mir-598-3p attenuated SEFM in SS, but not SR, males (F_(1,225)_ = 16.879, *p* < 0.001; LSD post hoc comparisons, SS/INH vs. SS/CTL, *p* < 0.05 for all comparisons, SR/INH vs. SR/CTL, *p* > 0.05 for all comparisons), indicating that mir-598-3p inhibition interferes with remote memory, but only when mice display stress-induced enhancement. Importantly, this interaction is not due to a floor effect, as mir-598-3p inhibition attenuated SEFM in SS, but not SR, even at Bins 2 and 3, before SR mice had extinguished and when normalized freezing is still ∼ 20%. Consistent with this, mir-598-3p inhibition increased the rate of extinction learning in SS mice only (two way ANOVA, treatment by population interaction F_(1,44)_ =4.767, *p* = 0.034; LSD posthoc comparisons SS/CTL vs. SS/INH, *p* = 0.021; SR/CTL vs. SR/INH, *p* = 0.424).

BLA mir-598-3p inhibition had no effect on SEFM in female mice (**Figure 2F**; two-way RM ANOVA, no main effect of treatment: F_(1,65)_ = 0.098, *p* = 0.755), consistent with the lack of a SEFL-induced increase in mir-598-3p levels (**Figure 2B**). In addition, freezing behavior did not decrease across the extinction session (no main effect of CS:, F_(4,65)_ = 1.91, *p* = 0.119), consistent with our previous report of a uniform memory enhancement displayed by females in response to stress [25].

### Amygdala mir-598-3p does not alter stress-induced changes in anxiety-like behavior in SEFL or FC mice

A potential interpretation of the reduced SEFM by intra-BLA mir-598-3p inhibition (**Figure 2E**) is that it produced an anxiolytic-like effect. This is plausible, given that BLA mir-598-3p is increased following both SEFL and restraint stress alone. Therefore, we next tested the effect of in vivo manipulation of mir-598-3p on anxiety-like behaviors observed in male mice exposed to SEFL or FC alone. Twenty-eight days after SEFL or FC, mice received intra-BLA infusions of mir-598-3p-INH or a nontargeting control. Anxiety-like behavior was then measured in the elevated plus maze (EPM), open field test (OFT), and acoustic startle response test (ASR) over three consecutive days (**Figure 3A**). In FC only mice, mir-598-3p inhibition in the same FC only mice appeared to produce an anxiolytic-like phenotype in the OFT (**Figure 3B**; t_(21)_ = 2.305, *p* = 0.032) without altering total distance travelled (**Figure 3C**; t_(21)_ = 1.361, *p* = 0.188). However, mir-598-3p inhibition did not alter time in the open versus closed arm of the EPM (**Figure 3D**; t_(20)_ = 0.571, *p* = 0.574), nor was there a change in total distance travelled in the maze (**Figure 3E**; t_(20)_ = 1.020, *p* = 0.312). There was no effect of mir-598-3p inhibition on acoustic startle in FC only mice (**Figure 3F**; t_(22)_ = 1.736, *p* = 0.097).

**Figure 3.**
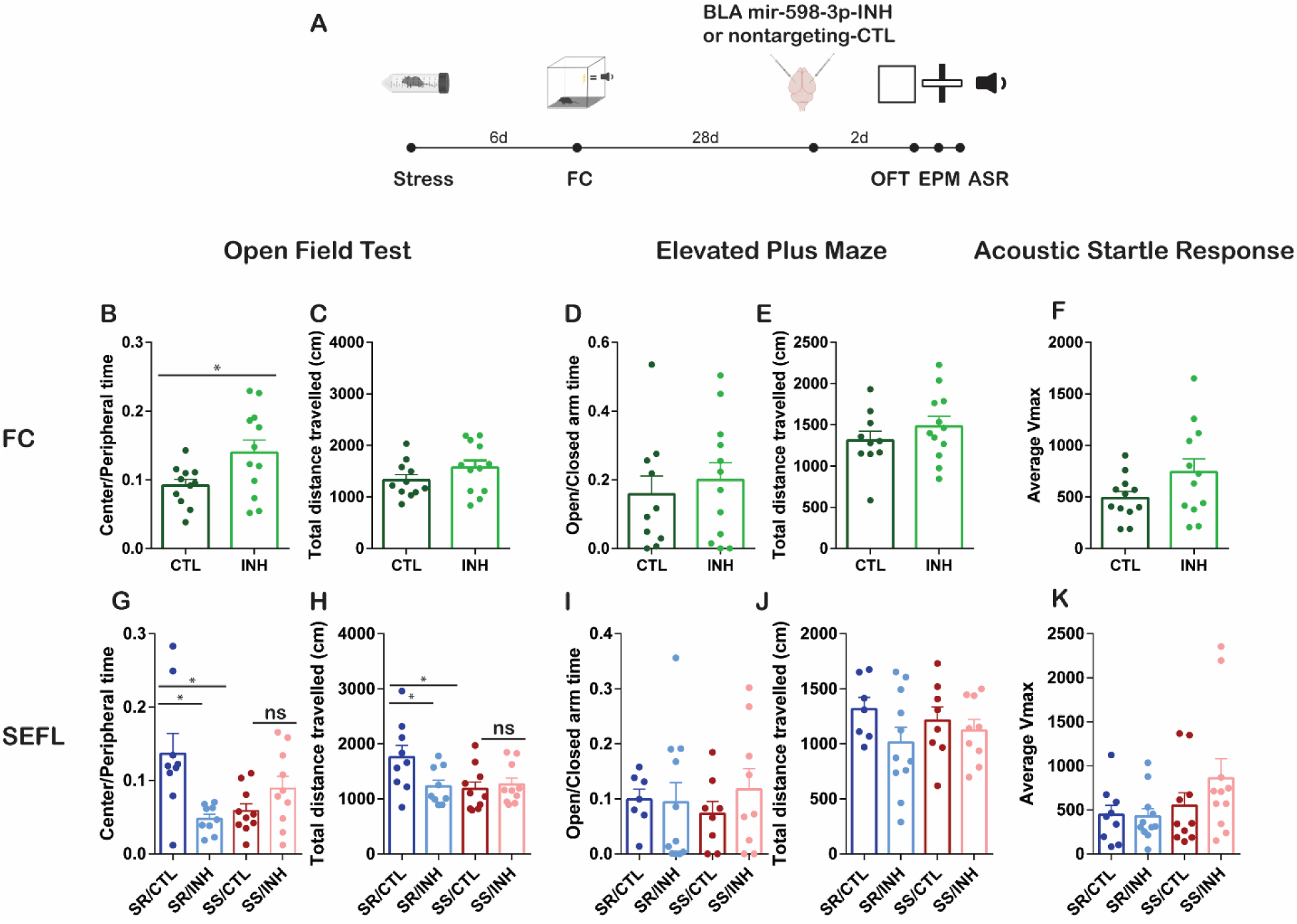
Amygdala mir-598-3p inhibition does not alter anxiety-like behavior in SEFL mice. (A) Schematic of experimental design. Center/Peripheral time (A) and total distance travelled (B) in the open field test (OFT) are presented in FC only mice. Open/Closed time (D) and total distance travelled (E) in the elevated plus maze (EPM) are presented in FC only mice. Average Vmax during the first 5 tones of the acoustic startle response (ASR) test (F) are presented in FC only mice. Center/Peripheral time (G) and total distance travelled (H) in the OFT are presented in SEFL mice. Open/Close time (I) and total distance travelled (J) in the EPM are presented in SEFL mice. Average Vmax during the first 5 tones of the ASR test (K) are presented in SEFL mice. * *p* < 0.05; ns = not significantly different; FC n = 10-12 per group; SR n = 7-11 per group, SS n = 8-10 per group.

Thirty days after SEFL training, we observed a significant population by mir-598-3p treatment interaction on anxiety-like behavior in the OFT (**Figure 3G**; F_(1,34)_ = 12.657, *p* = 0.001). Post hoc analyses confirmed an anxiogenic-like phenotype in SS mice 30 days post-SEFL, as compared to SR controls (SS/CTL vs. SR/CTL, *p* = 0.002), that was not rescued by mir-598-3p treatment (SS/CTL vs. SS/INH, *p* = 0.194). In addition, mir-598-3p inhibition was anxiogenic-like in SR mice (SR/CTL vs. SR/INH, *p* = 0.001), in which inhibition has no effect on SEFM. There was also a significant interaction effect in the total distance travelled in the OFT (**Figure 3H**; F_(1,34)_ = 4.407, *p* = 0.043; SS/CTL vs. SR/CTL, *p* = 0.008; SS/CTL vs. SS/INH, *p* = 0.684; SR/CTL vs. SR/INH, *p* = 0.017). Importantly, mir-598-3p inhibition did not affect anxiety-like behavior or distance travelled in the OFT in SS Mice. This was unexpected in that SS inhibitor mice display reduced fear memory on par with SR controls, but these data suggest that mir-598-3p inhibition does not similarly protect against any stress-induced increases in anxiety-like behavior in the open field. Taken together, these results suggest a decoupling of stress from anxiety-like behaviors. mir-598-3p inhibition produced no change in anxiety-like behavior in the EPM (**Figure 3I**; F_(1,31)_ = 0.362, *p* = 0.552) and there was no effect of population between SS and SR control mice (**Figure 3I**; F_(1,31)_ = 0.002, p = 0.969). There was also no effect of mir-598-3p-INH (**Figure 3J**; F_(1,31)_ = 2.464, *p* = 0.127) or of population (**Figure JH**, F_(1,31)_ = 0.001, *p* = 0.970) on the total distance travelled in the maze. Similarly, there was no effect of mir-598-3p (**Figure 3K**; F_(1, 37)_ = 0.913, *p* = 0.346) or population (**Figure 3K**; F_(1,37)_ = 3.006, p = 0.091) on acoustic startle response in SR or SS mice. Collectively, these data suggest that an anxiogenic-like phenotype in the open field test develops in SS mice over time, as thigmotaxis is increased and general locomotor exploration decreased in control animals 30 days post-SEFL, but not at the earlier time point of 4 days post-SEFL we previously assessed [25]. Importantly, mir-598-3p is not functionally related to this effect or to anxiety-like behavior in the EPM or ASR.

### Putative mir-598-3p targets are differentially expressed in the BLA of stress susceptible and resilient mice

miRNAs have the potential to titrate expression levels of multiple protein targets, creating complex miRNA-mRNA interaction patterns that may underlie functional consequences of the in vivo manipulations of mir-598-3p described here. We examined the BLA proteome in SS and SR mice 30 days after SEFL training (Sillivan et al, resubmitted; **Figure 4A**) and used three miRNA databases that organize known miRNA sequences and predict protein targets based on seed sequences to identify potential functional targets of mir-598-3p (DIANA [26], TargetScan [27], and miRbase [28]). Proteins predicted by at least two of three databases were considered to be potential targets of mir-598-3p. Twenty-three percent of predicted mir-598-3p targets were detected in the amygdala at the level of the proteome (**Figure 4B**). Proteins that differed by at least 0.5 log_2_ fold change were considered to be differentially expressed between SS and SR mice and of these, 29% were downregulated in SS mice, 45% were upregulated in SS mice, and 26% were not differentially expressed (**Figure 4C**). The 10 proteins that are predicted targets of mir-598-3p and are differentially expressed in SS compared to SR mice to the greatest degree, both up and down, are presented in **Figure 4D**. Because mir-598-3p is upregulated in SS mice, proteins functionally related to the protective effect of mir-598-3p inhibition on SEFL would be expected to be downregulated in SS mice. To explore potential functional annotations related to our observed effect, we conducted an Ingenuity Pathway Analysis on these downregulated putative mir-598-3p targets and identified several functional pathways that are involved in synaptic transmission and synaptic plasticity, including “density of excitatory synapses” and “quantity of dendritic spines”. Interestingly, several pathways point specifically to the regulation of the actin cytoskeleton, a crucial component of structural plasticity supporting functional plasticity [29-31]. Selected functional annotations and the protein targets associated with each are presented in **Figure 4E**. This includes representation of Pak3 in several functional annotations, which interacts directly with Rho GTPases to regulate the actin cytoskeleton [32].

**Figure 4.**
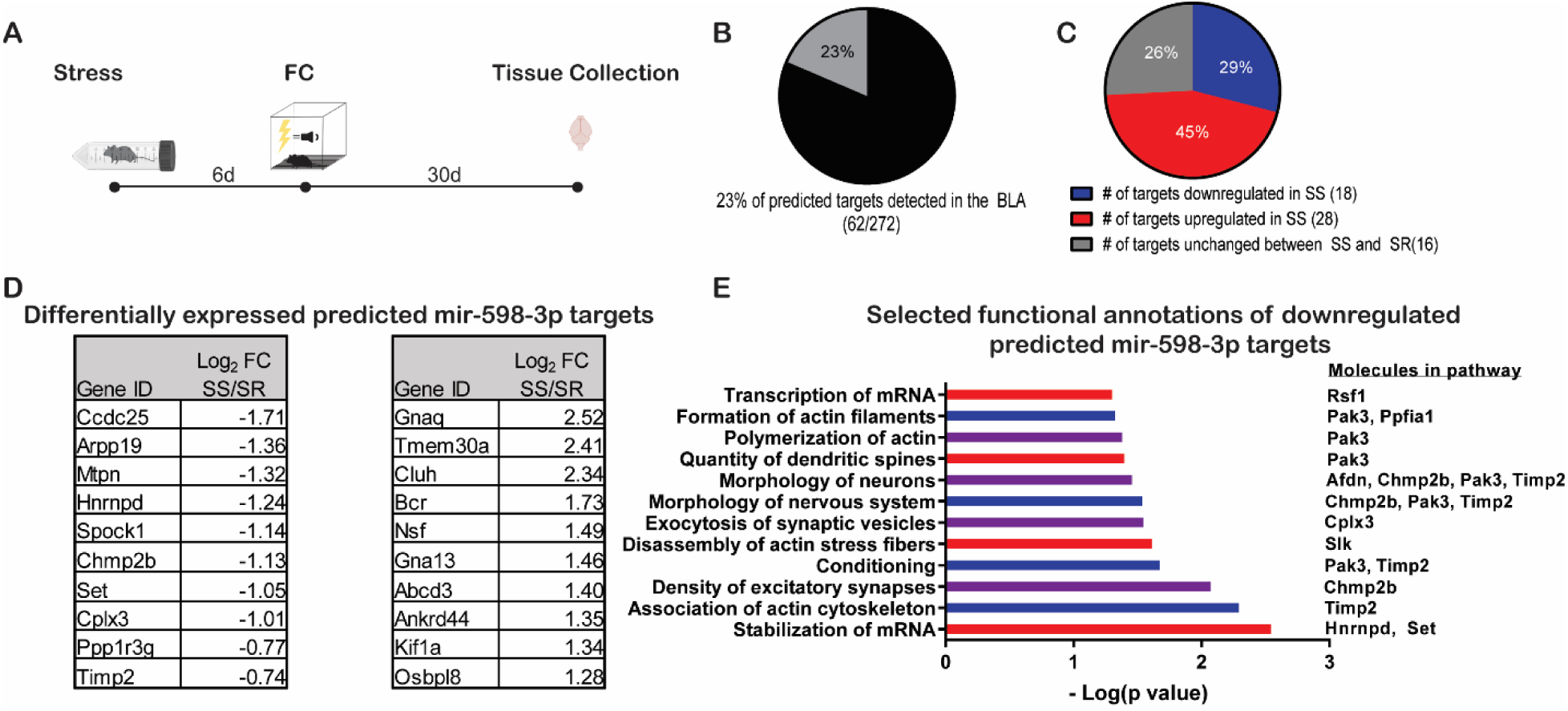
Putative mir-598-3p targets are differentially expressed in the BLA of stress susceptible and resilient mice. (A) Schematic of experimental design. (B) 272 protein targets were predicted by at least 2 of the three databases, 23% of those were detected at the level of the proteome in the BLA. (C) 29% of predicted targets were downregulated in SS mice, 45% of predicted targets were upregulated in SS mice, and 26% of predicted targets were not differentially expressed between SS and SR mice. (D) Log_2_ fold change (FC) values from top regulated predicted protein targets of mir-598-3p are presented. (E) Selected functional annotations from an Ingenuity Pathway Analysis of predicted mir-598-3p targets that were downregulated in SS mice compared to SR and the molecules associated with each.

## DISCUSSION

The smRNA-seq results presented here identify nine BLA miRNAs downregulated one month after cued fear conditioning, suggesting they may contribute to the persistence of remote memory. Interestingly, when assessing remote time points (one month post-training), we found that mir-598-3p is downregulated in the BLA following cued FC, but upregulated after exposure to acute stress. While it does not preclude the possibility that overexpressing mir-598-3p in the BLA would disrupt a remote fear memory, we found that further reduction of its levels in the BLA did not produce memory enhancement. Subsequently, we employed our SEFL protocol [25], in which mice are exposed to an acute stressor followed by cued FC one week later, to examine the role of mir-598-3p in susceptibility and/or resilience to the development of a remote SEFM. Thirty days after SEFL, amygdala mir-598-3p was increased in SS compared to SR mice and in vivo inhibition of mir-598-3p attenuated SEFM in SS mice, mimicking the SR phenotype. Further, we observed an anxiety-like phenotype in SS mice 30 days after SEFL relative to SR mice that was not altered by the same manipulation of mir-598-3p. Thus, the effect of BLA mir-598-3p inhibition was specific to memory, failing to influence general anxiety-like behaviors. Consistent with a memory-specific role for mir-598-3p, pathway analysis of putative targets of mir-598-3p that were found to be downregulated in SS mice compared to SR in our quantitative proteomics dataset revealed several functional pathways related to plasticity underlying learning and memory.

In the first description of a behavioral effect linked to mir-598-3p in the brain, we show SS mice fail to exhibit the time-dependent reduction in BLA mir-598-3p levels that FC and SR groups exhibited 30 days after learning. BLA mir-598-3p inhibition in SS mice protected against SEFM and restored extinction learning to SR and FC levels, without a comparable anxiolytic effect. The memory-, SS- and sex-specific effect of mir-598-3p manipulation suggests that the functional pathways targeted are specifically related to the impact of stress on memory. Interestingly, a precursor miRNA family, mir-19b, has been implicated in PTSD [20, 21] and is differentially expressed between SS and SR mice in our own laboratory. This same miRNA is enhanced in the amygdala following chronic social defeat stress and mir-19b in vivo manipulation bidirectionally modulated cued fear conditioning without altering contextual fear memory or anxiety-like behavior in the OFT, dark/light transfer test, or the EPM [14]. Further study of miRNAs altered by stress, such as mir-19b or mir-598-3p, which specifically influence amygdala-dependent memory without altering general anxiety-like behavior, may provide further insight into mechanisms contributing to PTSD.

The pathway analysis conducted on putative targets of mir-598-3p detected to be downregulated in SS mice revealed several pathways linked to learning and plasticity. Plasticity of dendritic spines is critical for long-term memory formation [33] and is supported by actin polymerization [34]. Several functional annotations we report here that are related to actin dynamics involved two proteins in particular, P21 (Rac1) activated kinase 3 (Pak3) and tissue inhibitor of metallaoproteinase 2 (Timp2). Pak3 interacts directly with Rho GTPases to regulate actin dynamics driving rapid cytoskeletal reorganization in the adult brain [32]. It is most commonly associated with intellectual disability [35] and inactivation of Pak3 in the forebrain is associated with impaired spatial memory in the Morris Water Maze and contextual fear conditioning, while sparing cued fear conditioning memory [36]. This is interesting, given that our results suggest lower levels of Pak3 are associated with enhanced memory following a stressor, rather than impaired memory, and suggests that a background of stress might alter endogenous levels of Pak3. Timp2 regulates degradation of the extracellular matrix [37]. While there is little known about the direct role of Timp2 in learning and memory, the extracellular matrix maintains direct contact with synapses and is known to influence dendritic spine stability and synapse structure [38]. In addition, it is reduced in patients with recurrent depressive symptoms [39]. The potential connection to depressive symptoms is interesting given the downregulation observed in our PTSD-like phenotype (SS mice) and the high comorbidity between PTSD and major depressive disorder in patients [40].

While there are no reports of the effect of central mir-598-3p on learning in the literature, mir-598-3p is downregulated in the cerebrospinal fluid of patients diagnosed with Alzheimer’s disease [41]. Given that Alzheimer’s is associated with significant memory impairment as the disease progresses, this is consistent with upregulated mir-598-3p in SS mice without retrieval being associated with an enhanced fear memory. While there was no effect of amygdala mir-598-3p inhibition in FC only or SR mice, the effect of mir-598-3p on memory could be target-dependent, in that baseline levels of mir-598-3p and its putative targets could determine which functional targets the miRNA interacts with in different contexts. This is consistent with the distinct anxiety-like effects of mir-598-3p inhibition in the OFT that we observed between FC, SR and SS treatment groups, that did not correspond to levels of mir-598-3p at the time of inhibition.

We observed opposing effects of mir-598-3p inhibition in FC (anxiolytic-like) and SR (anxiogenic-like) mice in the OFT. Interestingly, Dicer1 knockdown [23] and miR-135a inhibition [42, 43] in the amygdala were anxiogenic-like in the EPM. In the absence of a background of stress, miRNAs have been hypothesized to influence anxiety-like behavior potentially through metabolic or inflammatory mechanisms as several miRNAs, for example mir-34c and mir-26a family miRNAs, are implicated in both changes in inflammation and the modulation of anxiety-like behavior [23, 24, 44]. Inflammation is traditionally associated with enhanced anxiety [45], and consistently, mir26a has been shown to be anti-inflammatory and anxiolytic [24, 46]. Future studies could explore the role of other putative targets of mir-598-3p involved in inflammation to better understand its potential role in anxiety in either naïve or stress-resilient mice.

In summary, we identified amygdala mir-598-3p as one miRNA supporting the persistence of memory that is regulated by stress and may underlie remote stress-enhanced fear memory. Our findings suggest the effect of mir-598-3p inhibition is specific to susceptible mice and to memory, as there was no anxiolytic affect in the EPM, OFT, or ASR associated with the effect. Further study of putative targets of mir-598-3p in SS mice could ultimately provide insight for treatments aimed at targeting the selective disruption of traumatic memory.

## MATERIALS AND METHODS

### Animals

Adult C57BL/6 mice, 8 weeks of age (The Jackson Laboratory, Bar Harbor, ME), were maintained on a 12:12 hour light/dark cycle and supplied with food and water ad libitum. Animals were housed 3-4/cage, acclimated to the facility for 1 week then handled for 3 days prior to experiments. Behavioral tests were performed between 8AM and 5PM. Treatment groups were randomized for all behavioral experiments to prevent batch effects due to time of day. Procedures were performed in accordance with the Institutional Animal Care and Use Committee (IACUC) at the Scripps Florida Research Institute and with national regulations and policies.

### Behavioral paradigms

Stress-enhanced fear learning (SEFL) was performed as previously described [25]. The procedure combines a single acute restraint stress session with auditory fear conditioning to produce stress susceptible (SS) and stress resilient (SR) populations of animals. Freezing during the last 60 seconds of shock training was used to determine SS/SR classifications, as this measure correlates with long-lasting extinction resistance and fear memory expression. Extinction with 30 tone presentations was performed in a context unique from training. All freezing behavior was normalized to pretone freezing to isolate cue-specific freezing and the rate of habituation was calculated as ((average % freezing during the first three tones) – (average % freezing during the last three tones) / (average % freezing during the first three tones) * 100). Open field, elevated plus maze, and acoustic startle response tests were performed as previously described [25, 47]. Behavior in the elevated plus maze was analyzed as open arm time/ closed arm time and behavior in the open field test was analyzed as time spent in the center/time spent in the periphery of the maze. For both tests, behavior was analyzed for a total of 5 minutes and total distance travelled during the test was also analyzed. For acoustic startle, mean startle amplitude for the first 5 stimuli was analyzed. For traditional fear conditioning, SEFL and anxiety battery experiments, mice underwent stress and/or fear conditioning and then remained in the home cage prior to surgery for intra-amygdalar miRNA inhibitor surgery.

### RNA extraction and qPCR

Total RNA from fresh frozen bilateral tissue punches was obtained using the miRVANA PARIS RNA extraction kit (Life Technologies, Carlsbad, CA), as previously reported [5, 48]. cDNA libraries of miRNAs were created from 20ng of total RNA with the mirCURY LNA RT Kit (Qiagen, Germantown, MD). PCR reactions were performed using the miRCURY LNA SYBR Green PCR Kit and the following locked nucleic acid (LNA) SYBR green primers from Qiagen: mmu-miR-598-3p, assay ID: YP00205045; snord68, assay ID: YP00203911; and RNU5G, assay ID: YP00203908. Data were normalized to the housekeeping genes snord68 and RNU5G using the ΔΔct method [49].

### Quantitative mass spectrometry

Mass spectrometry was performed at the Harvard Mass Spectrometry and Proteomics Resource Laboratory as previously described [5] and data was mined from a quantitative mass spectrometry run on BLA tissue collected from male mice 30 days post-SELF training (reported in Sillivan et al, *resubmitted*). Candidate proteins were identified as those that had at least 0.5 log2 fold change between treatment groups.

### Intra-amygdalar infusions

Hairpin inhibitors directed against mmu-miR-598-3p or the nonmammalian miRNA cel-miR-67 were obtained from GE Dharmacon (Lafayette, CO) and injected bilaterally into the BLA (AP: 1.5 mm, ML: ±3.2 mm from bregma and DV: – 4.7 mm from the skull) as previously described [50]. The injection needle was left in place for 5 minutes after the infusion was complete. Inhibitors were reconstituted in water then prepared with jetPEI transfection reagent (Polyplus Transfection, Illkirch, France). 1ul of 400ng/ul was injected 28 days after fear conditioning, animals remained in the home age for two days, and then were tested for either remote fear memory expression or anxiety-like behavior.

### Dual fluorescence in situ hybridization (FISH)

Neuroanatomical localization of mir-598-3p was achieved with FISH. Expression analysis experiments to detect miRNAs were performed as described (reported in Sillivan et al, *resubmitted*) using LNA dual 5’- and 3’-DIG-labeled probes against mir-598-3p or a scrambled negative control sequence (Exiqon). Sections were mounted with Prolong Gold Diamond antifade mounting media with DAPI (Thermo Fisher) and visualized on an Olympus fluorescent confocal microscope with 60X objective (Tokyo, Japan).

### Cell fractionation

BLC tissue from naïve animals was dissected fresh and immediately separated into cytosol/nuclear and synaptosome fractions as previously described (reported in Sillivan et al, *resubmitted*). Each sample was pooled from four animals. RNA was extracted from each cellular fractions with the miRVANA PARIS kit as described above.

### Statistical Analysis

A student’s t test was used to analyze the initial qPCR experiment validating the FC-induced reduction relative to naïve. A one-way ANOVA was used to analyze mir-598-3p expression in each of the 4 (for female) or 5 (for male) groups analyzed at 30 days, 7 days, or 24 hours after each behavioral manipulation. Two-way (FC only) or three-way (SEFL) repeated measures ANOVAs were used to analyze the effect of mir-598-3p inhibition on remote fear memory in FC mice and SEFL mice, respectively. Significant interactions were reported and analyzed and with LSD posthoc tests were used for pairwise comparisons. Student’s t tests (FC only) and two-way ANOVAs (SEFL) were used to analyze anxiety-like behavior in the EPM, OFT, and ASR tests, respectively. LSD posthoc tests were used for pairwise comparisons.

## ACKNOWLEDGEMENTS

We thank the Scripps Florida Genomics Core for sequencing services, Nripesh Prasad at the Genomic Services Lab at Hudson Alpha for sequencing services and data analysis, Adrian Reich and the Bioinformatics Core for data analysis, the Mouse Behavior core and Alicia Brantley for assistance and behavioral equipment, all members of the Miller and Rumbaugh Labs for their technical assistance and thoughtful discussions. This work was funded by grants from the National Institute of Mental Health MH105400 (CM), National Institute on Drug Abuse DA041469 (SS) and the Brain and Behavior Foundation-NARSAD Young Investigator Award (SS).

## REFERENCES

1. Gisquet-Verrier, P. and C. Le Dorze, Post Traumatic Stress Disorder and Substance Use Disorder as Two Pathologies Affecting Memory Reactivation: Implications for New Therapeutic Approaches. Front Behav Neurosci, 2019. 13: p. 26.

2. Bowers, M.E. and K.J. Ressler, An Overview of Translationally Informed Treatments for Posttraumatic Stress Disorder: Animal Models of Pavlovian Fear Conditioning to Human Clinical Trials. Biol Psychiatry, 2015. 78(5): p. E15–27.

3. Asok, A., et al., Molecular Mechanisms of the Memory Trace. Trends Neurosci, 2019. 42(1): p. 14–22.

4. Brewin, C.R., Re-experiencing traumatic events in PTSD: new avenues in research on intrusive memories and flashbacks. Eur J Psychotraumatol, 2015. 6: p. 27180.

5. Sillivan, S.E., et al., Bioinformatic analysis of long-lasting transcriptional and translational changes in the basolateral amygdala following acute stress. PLoS One, 2019. 14(1): p. e0209846.

6. McEwen, B.S., Physiology and neurobiology of stress and adaptation: central role of the brain. Physiol Rev, 2007. 87(3): p. 873–904.

7. Zannas, A.S. and A.E. West, Epigenetics and the regulation of stress vulnerability and resilience. Neuroscience, 2014. 264: p. 157–70.

8. Gebert, L.F.R. and I.J. MacRae, Regulation of microRNA function in animals. Nat Rev Mol Cell Biol, 2019. 20(1): p. 21–37.

9. Filipowicz, W., S.N. Bhattacharyya, and N. Sonenberg, Mechanisms of post-transcriptional regulation by microRNAs: are the answers in sight? Nat Rev Genet, 2008. 9(2): p. 102–14.

10. Konopka, W., et al., MicroRNA loss enhances learning and memory in mice. J Neurosci, 2010. 30(44): p. 14835–42.

11. Gao, J., et al., A novel pathway regulates memory and plasticity via SIRT1 and miR-134. Nature, 2010. 466(7310): p. 1105–9.

12. Griggs, E.M., et al., MicroRNA-182 regulates amygdala-dependent memory formation. J Neurosci, 2013. 33(4): p. 1734–40.

13. Dias, B.G., et al., Amygdala-dependent fear memory consolidation via miR-34a and Notch signaling. Neuron, 2014. 83(4): p. 906–18.

14. Volk, N., et al., MicroRNA-19b associates with Ago2 in the amygdala following chronic stress and regulates the adrenergic receptor beta 1. J Neurosci, 2014. 34(45): p. 15070–82.

15. Lin, Q., et al., The brain-specific microRNA miR-128b regulates the formation of fear-extinction memory. Nat Neurosci, 2011. 14(9): p. 1115–7.

16. Gale, G.D., et al., Role of the basolateral amygdala in the storage of fear memories across the adult lifetime of rats. J Neurosci, 2004. 24(15): p. 3810–5.

17. Mantzur, L., G. Joels, and R. Lamprecht, Actin polymerization in lateral amygdala is essential for fear memory formation. Neurobiol Learn Mem, 2009. 91(1): p. 85–8.

18. Zelikowsky, M., et al., Neuronal ensembles in amygdala, hippocampus, and prefrontal cortex track differential components of contextual fear. J Neurosci, 2014. 34(25): p. 8462–6.

19. Rodrigues, S.M., G.E. Schafe, and J.E. LeDoux, Molecular mechanisms underlying emotional learning and memory in the lateral amygdala. Neuron, 2004. 44(1): p. 75–91.

20. Martin, C.G., et al., Circulating miRNA associated with posttraumatic stress disorder in a cohort of military combat veterans. Psychiatry Res, 2017. 251: p. 261–265.

21. Zhou, J., et al., Dysregulation in microRNA expression is associated with alterations in immune functions in combat veterans with post-traumatic stress disorder. PLoS One, 2014. 9(4): p. e94075.

22. Cohen, J.L., et al., Differential stress induced c-Fos expression and identification of region-specific miRNA-mRNA networks in the dorsal raphe and amygdala of high-responder/low-responder rats. Behav Brain Res, 2017. 319: p. 110–123.

23. Haramati, S., et al., MicroRNA as repressors of stress-induced anxiety: the case of amygdalar miR-34. J Neurosci, 2011. 31(40): p. 14191–203.

24. Xie, L., et al., MicroRNA-26a-2 maintains stress resiliency and antidepressant efficacy by targeting the serotonergic autoreceptor HTR1A. Biochem Biophys Res Commun, 2019. 511(2): p. 440–446.

25. Sillivan, S.E., et al., Susceptibility and Resilience to Posttraumatic Stress Disorder-like Behaviors in Inbred Mice. Biol Psychiatry, 2017. 82(12): p. 924–933.

26. Karagkouni, D., et al., DIANA-TarBase v8: a decade-long collection of experimentally supported miRNA-gene interactions. Nucleic Acids Res, 2018. 46(D1): p. D239–D245.

27. Agarwal, V., et al., Predicting effective microRNA target sites in mammalian mRNAs. Elife, 2015. 4.

28. Kozomara, A., M. Birgaoanu, and S. Griffiths-Jones, miRBase: from microRNA sequences to function. Nucleic Acids Res, 2019. 47(D1): p. D155–D162.

29. Hotulainen, P. and C.C. Hoogenraad, Actin in dendritic spines: connecting dynamics to function. J Cell Biol, 2010. 189(4): p. 619–29.

30. Kasai, H., et al., Structural dynamics of dendritic spines in memory and cognition. Trends Neurosci, 2010. 33(3): p. 121–9.

31. Novaes, L.S., et al., Environmental enrichment prevents acute restraint stress-induced anxiety-related behavior but not changes in basolateral amygdala spine density. Psychoneuroendocrinology, 2018. 98: p. 6–10.

32. Ramakers, G.J., Rho proteins, mental retardation and the cellular basis of cognition. Trends Neurosci, 2002. 25(4): p. 191–9.

33. Yang, G., F. Pan, and W.B. Gan, Stably maintained dendritic spines are associated with lifelong memories. Nature, 2009. 462(7275): p. 920–4.

34. Smart, F.M. and S. Halpain, Regulation of dendritic spine stability. Hippocampus, 2000. 10(5): p. 542–54.

35. Muthusamy, B., et al., Next-Generation Sequencing Reveals Novel Mutations in X-linked Intellectual Disability. OMICS, 2017. 21(5): p. 295–303.

36. Hayashi, M.L., et al., Altered cortical synaptic morphology and impaired memory consolidation in forebrain-specific dominant-negative PAK transgenic mice. Neuron, 2004. 42(5): p. 773–87.

37. Perez-Martinez, L. and D.M. Jaworski, Tissue inhibitor of metalloproteinase-2 promotes neuronal differentiation by acting as an anti-mitogenic signal. J Neurosci, 2005. 25(20): p. 4917–29.

38. Levy, A.D., M.H. Omar, and A.J. Koleske, Extracellular matrix control of dendritic spine and synapse structure and plasticity in adulthood. Front Neuroanat, 2014. 8: p. 116.

39. Bobinska, K., et al., The role of MMP genes in recurrent depressive disorders and cognitive functions. Acta Neuropsychiatr, 2016. 28(4): p. 221–31.

40. Jaksic, N., B.A. Margetic, and D. Marcinko, Comorbid Depression and Suicide Ideation in Patients with Combat-Related PTSD: The Role of Temperament, Character, and Trait Impulsivity. Psychiatr Danub, 2017. 29(1): p. 51–59.

41. Riancho, J., et al., MicroRNA Profile in Patients with Alzheimer’s Disease: Analysis of miR-9-5p and miR-598 in Raw and Exosome Enriched Cerebrospinal Fluid Samples. J Alzheimers Dis, 2017. 57(2): p. 483– 491.

42. Issler, O., et al., MicroRNA 135 is essential for chronic stress resiliency, antidepressant efficacy, and intact serotonergic activity. Neuron, 2014. 83(2): p. 344–360.

43. Mannironi, C., et al., miR-135a Regulates Synaptic Transmission and Anxiety-Like Behavior in Amygdala. Mol Neurobiol, 2018. 55(4): p. 3301–3315.

44. Meydan, C., S. Shenhar-Tsarfaty, and H. Soreq, MicroRNA Regulators of Anxiety and Metabolic Disorders. Trends Mol Med, 2016. 22(9): p. 798–812.

45. Michopoulos, V., et al., Inflammation in Fear- and Anxiety-Based Disorders: PTSD, GAD, and Beyond. Neuropsychopharmacology, 2017. 42(1): p. 254–270.

46. Zhang, Y., et al., Effects of miR-26a-5p on neuropathic pain development by targeting MAPK6 in in CCI rat models. Biomed Pharmacother, 2018. 107: p. 644–649.

47. Ozkan, E.D., et al., Reduced cognition in Syngap1 mutants is caused by isolated damage within developing forebrain excitatory neurons. Neuron, 2014. 82(6): p. 1317–33.

48. Rumbaugh, G., et al., Pharmacological Selectivity Within Class I Histone Deacetylases Predicts Effects on Synaptic Function and Memory Rescue. Neuropsychopharmacology, 2015. 40(10): p. 2307–16.

49. Livak, K.J. and T.D. Schmittgen, Analysis of relative gene expression data using real-time quantitative PCR and the 2(-Delta Delta C(T)) Method. Methods, 2001. 25(4): p. 402–8.

50. Young, E.J., et al., Nonmuscle myosin IIB as a therapeutic target for the prevention of relapse to methamphetamine use. Mol Psychiatry, 2016. 21(5): p. 615–23.

